# Establishment of CD1b-restricted immunity to lipid antigens in the pulmonary response to *Mycobacterium tuberculosis* infection

**DOI:** 10.1101/2023.05.23.541963

**Authors:** Macallister C. Harris, Hadley E. Gary, Sarah K. Cooper, David F. Ackart, James E. Dilisio, Randall J Basaraba, Tan-Yun Cheng, Ildiko van Rhijn, D. Branch Moody, Brendan K. Podell

## Abstract

CD1 is an antigen presenting glycoprotein homologous to MHC I; however, CD1 proteins present lipid rather than peptide antigen. CD1 proteins are well established to present lipid antigens of *Mycobacterium tuberculosis* (Mtb) to T cells, but understanding the role of CD1-restricted immunity *in vivo* in response to Mtb infection has been limited by availability of animal models naturally expressing the CD1 proteins implicated in human response: CD1a, CD1b and CD1c. Guinea pigs, in contrast to other rodent models, express four CD1b orthologs, and here we utilize the guinea pig to establish the kinetics of gene and protein expression of CD1b orthologs, as well as the Mtb lipid-antigen and CD1b-restricted immune response at the tissue level over the course of Mtb infection. Our results indicate transient upregulation of CD1b expression during the effector phase of adaptive immunity that wanes with disease chronicity. Gene expression indicates that upregulation of CD1b is the result of transcriptional induction across all CD1b orthologs. We show high CD1b3 expression on B cells, and identify CD1b3 as the predominant CD1b ortholog in pulmonary granuloma lesions. We identify *ex vivo* cytotoxic activity directed against CD1b that closely paralleled the kinetic changes in CD1b expression in Mtb infected lung and spleen. This study confirms that CD1b expression is modulated by Mtb infection in lung and spleen, leading to pulmonary and extrapulmonary CD1b-restricted immunity as a component of the antigen-specific response to Mtb infection.

## Introduction

Expanding the perspective that T cells recognize MHC-peptide complexes, CD1-lipid complexes activate invariant natural killer T cells that specifically respond to CD1d, while other T cells recognize lipid antigens presented by CD1a, CD1b and CD1c (1, 2). CD1 glycoproteins have homologous structuring to MHC I antigen-presenting molecules but bind and present a variety of lipid antigens (3, 4). In humans, four CD1 antigen presenting molecules are divided into two functional groups; group 1 is composed of CD1a, CD1b and CD1c, and group 2 is composed of CD1d (5–7). Group 2 CD1 complexes are constitutively expressed on many antigen-presenting cell (APC) types, including macrophages, B cells and myeloid dendritic cells, where they present sphingoglycolipids for expansion of invariant natural killer T cells (8, 9). In contrast, Group 1 CD1 proteins have inducible expression on myeloid dendritic cells, where they load endogenous and exogenous lipids, such as those derived from *Mycobacterium tuberculosis* (Mtb), leading to an expansion of a variety of functionally nonpolymorphic CD1-restricted αβ and γδ T cells (10–13).

Owing to the lipid-rich composition of Mtb, multiple studies have investigated the potential role of CD1-restricted immunity in tuberculosis (TB). Studies examining peripheral blood of Mtb-infected human subjects found increased expression of CD1 on myeloid cells and corresponding presence of CD1-restricted T cells (14–17). Other studies have taken additional strides in isolation of Mtb-lipid specific T cells via detection with lipid-loaded CD1 tetramers (18, 19). Though these studies lay a foundational understanding of CD1-restricted immunity during Mtb infection by identifying polyclonal T cell responses *ex vivo*, many gaps remain: whether CD1-restricted immunity contributes as a component of innate or adaptive branches of immunity *in vivo*, whether CD1 expression is modulated by Mtb infection *in vivo*, and whether CD1 plays a functional role in disease pathogenesis and granuloma formation in the tissue response to infection (6, 7, 16).

One major limitation to further understanding of CD1 lipid-restricted immunity in response to Mtb is the lack of a translatable animal model with naturally occurring group 1 CD1 expression. Experimentally tractable mouse models naturally express only CD1d. However, guinea pigs have a similar CD1 complex repertoire to humans, containing 4 orthologs of the group 1 CD1 protein and 3 orthologs of CD1c, as well as CD1d (20, 21). Previous studies have characterized this CD1 repertoire *in vivo* and *ex vivo* as well as in the face of administered lipids isolated from cultured Mtb, and BCG vaccination (20, 22, 23). However, there is no understanding of how Mtb infection impacts CD1 expression and the development of CD1-restricted T cell immunity over the course of TB disease in infected tissue. In this study, we utilized the guinea pig model of pulmonary TB and reagents specific to CD1b orthologs to kinetically measure CD1b-restricted immunity at innate, early adaptive, and late adaptive stages of TB within the lung and spleen. Overall, this work demonstrates that CD1 immunity actively responds to Mtb, as manifested by CD1b-mediated cytolytic responses to mycobacterial lipid antigens, indicating a pathogen-specific *in vivo* response to infection. Further, this study establishes the ability to assess CD1 immunity at the protein and molecular level within infected tissue utilizing a translatable small animal model.

## Material and Methods

### Animals

Two separate animal studies were performed to kinetically monitor CD1b expression in blood and tissue over the course of Mtb infection. Female outbred Dunkin-Hartley guinea pigs weighing 250-300 g were obtained from either Charles River Laboratories (Wilmington, MA) or Elm Hill Labs (Chelmsford, MA). In the first study, guinea pigs infected with Mtb were evaluated at 14-, 30- and 60-days post-infection (DPI) (n=6 per endpoint). In the second study, Mtb-infected guinea pigs at 14, 30 and 60 DPI (n=5 per endpoint) were compared to uninfected, naïve guinea pigs (n=3) at each necropsy endpoint to compare impact of infection over baseline CD1b expression. Tissues and cells from inbred strain 13 guinea pigs, a colony maintained at Colarado State University, were utilized for assay development, including cytotoxicity, CD1 expression via flow cytometric analysis, and validation of antibodies for immunohistochemistry on fresh frozen tissue. All animals were monitored via the Colorado State University (CSU) Lab Animal Resources staff and veterinarians and performed in accordance with protocols approved by the Institutional Animal Care and Use Committee protocol number 1401.

### CD1-specific reagents

To separately measure responses to the four CD1b orthologs in guinea pigs, we used 104C1 guinea pig fibroblast cell lines transfected with CD1b1, CD1b2, CD1b3 and CD1b4 (20). Specific CD1b ortholog-expressing cell lines were compared to a mock-transfectant containing only the empty pcDNA3.1 vector. Synthetic lipid antigens, glucose monomycolate and mycolic acid (24), were used to eliminate the possibility of co-purifying peptide antigens in preparations, which was essential to avoid MHC-restricted T cell responses. These lipid antigens were chosen as known CD1b ligands with demonstrated antigen-specific response in previous studies with antigen-specific IFNγ production or antigen-specific CD1b tetramer-based detection of cells in human peripheral blood (24–26). CD1 transfected 104C1 cells (CD1B1-4 and CD1C1-3), and anti-guinea pig CD1 mouse monoclonal antibodies allowed analysis of ortholog specificity (Fig. S1 and Table S1) (20).

### Infection and euthanasia

Low-dose aerosol exposure of guinea pigs to Mtb was performed using the Madison chamber aerosol generation device (College of Engineering Shops, University of Wisconsin, Madison, WI) calibrated to deliver approximately 20-50 bacilli of the H37Rv strain of Mtb isolated during log-phase growth in Proskauer-Beck media. A second study used the Glas-Col aerosol chamber (Terre Haute, IN) to deliver a low-dose exposure of <50 bacilli of the Erdman strain of Mtb. On pre-determined endpoints, 14, 30 and 60 days post-infection, guinea pig groups were euthanized via an initial induction of an anesthetic state via intramuscular injection of 30 mg of ketamine and 8 mg of xylazine in combination, followed by an intraperitoneal injection of 3 ml/animal of 390 mg/ml sodium pentobarbital.

### Necropsy and tissue processing

Animal tissues and cells were harvested, which included airway-accessible macrophages and other leukocytes by bronchoalveolar lavage, PBMCs from anticoagulated peripheral blood, and cell suspensions derived from lung and splenic parenchyma, as previously reported (27, 28). In brief, the right cranial and caudal bronchi were clamped and lung lobes removed and processed for either RNA preservation in RNAlater solution (Thermofisher, Waltham, MA), OCT-embedding for fresh-frozen immunohistochemistry (Sakura, Torrance, CA), or for bacterial enumeration by colony forming units (CFU). Airway-accessible leukocytes were collected via postmortem bronchioalveolar lavage utilizing an 18-gauge catheter and flushing with 25 ml of HBSS containing 0.01 μM EDTA solution in five, 5 ml, intervals. Following bronchoalveolar lavage, lung was perfused with 5 μg/ml collagenase (Sigma-Aldrich, cat#: C9263) and incubated for 30 minutes in a 37°C water bath. The digested lung parenchyma and collected spleen were individually processed through 40 μm cell strainers. Once strained, all samples were subjected to hypotonic solution for 20 to 45 seconds for lysis of erythrocytes. To increase overall cell viability for downstream assays, all tissue-derived cell suspensions were subjected to magnetic bead negative selection using biotinylated annexin V and streptavidin-conjugated magnetic nanobeads. Cells were placed in 4 ml of 1X annexin binding buffer (Biolegend, cat#: 422201) with biotinylated annexin V (Biolegend, Cat # 640904) and incubated for 30 minutes then centrifuged at 500 x g for 5 minutes, washed with binding buffer, centrifuged again and resuspended in 500 μl of binding buffer. Thirty μl of streptavidin nanobeads (Biolegend, cat#: 480016) were incubated for 20 minutes with intermittent vortexing. Additional binding buffer was added to bring solutions to 4 ml and the cell suspension was placed on a magnetic rack for 10 minutes to allow for magnetic binding of nanobeads. Finally, unbound cells were collected with a serologic pipette. Collected cells were washed in Hank’s balanced salt solution then viability and cell count assessed via a hemocytometer and trypan blue exclusion. Splenic and lung suspension cells were used for analysis of CD1b expression by flow cytometry and remaining cells plated in RPMI media containing 10% fetal bovine serum and an antimicrobial cocktail, then incubated overnight at 37°C with 5% CO_2_ to allow for separation of adherent cells (histiocytic/dendritic in origin) and non-adherent cells (lymphocytes). PBMCs were collected via a right ventricular intracardiac blood draw directly following euthanasia, diluted 1:2 in HBSS and separated on a density gradient using Lympholyte mammal media (Cedarlane, cat#: CL5110) per manufacturer’s instructions.

### Intradermal *M. tuberculosis* antigen challenge

In the second study, 48-hours prior to euthanasia, guinea pigs received an intradermal hypersensitivity challenge to either Mtb lipid antigens, Mtb protein antigen in the form of PPD, or sterile PBS. Each guinea pig received seven intradermal injections: 1. mycolic acid liposome, 2. glucose monomycolate liposome, 3. mycolic acid in PBS, 4. glucose monomycolate in PBS, 5. unloaded liposome, 6. Mtb purified protein derivative (PPD) (Tuberculin PPD Bovine, USDA Veterinary Services), or 7. sterile PBS. Liposomes were made using distearoylphosphatidylcholine (DSPC) (Avanti lipids, cat#: 850365P) and cholesterol (Avanti lipids, cat#: 700100P) lipids with or without synthetic Mtb lipid (molar ratio 7:2:1, respectively) via a lipid extruder (Avanti Lipids, Alabaster, AL). For liposomal formulation, DSPC and cholesterol were added to a pre-rinsed glass container, dissolved in a solution of 10:1 chloroform and methanol, and solubilized mycolic acid or glucose monomycolate added. The lipid solution was completely evaporated utilizing a nitrogen gas bath. Dried lipids were then suspended in PBS prewarmed to 68°C. The solution was then vortexed and incubated in a water bath at 68°C for 40 minutes, with vortexing every 10 minutes, to allow for liposome formation. During the incubation, the liposome extruder apparatus (Avanti lipids, Alabaster, AL) was assembled to company specifications and warmed to 70°C. After incubation, the liposome solution was passed through the extruding system 7-8 times to allow for formation of a uniform unilamellar liposomes at approximately 150 μm in diameter. Liposomal size was confirmed with a Nanosight instrument and capacity for macrophage phagocytosis confirmed using guinea pig bone marrow-derived macrophages in *in vitro* culture (Fig. S2). PBS lipid suspensions were sonicated using a Misonix ultrasonic liquid processor at 80 A for 4 minutes on a 20-second-on, 20-second-off cycle. All lipid injection sites received 4 μg of lipid whether in liposomal formulation or free lipid in PBS. PPD was administered at 2 μg per site. Guinea pigs were ventrally shaved from the point of the caudal xiphoid to the cranial aspect of the pubis. Injection sites were labelled and administered via intradermal injection using an insulin syringe at 50 μl of sample per site. All sites were measured at 24 hours and 48 hours after the initial injection using calipers to measure cranial to caudal and lateral diameters.

### Kinetic CD1 expression

CD1b expression was measured by flow cytometry on cell suspensions from lung and spleen, bronchoalveolar lavage, and PBMCs. In brief, published methods (27, 28) used 1.5 x 10^6^ cells that were blocked with 2.24 μg/ml of guinea pig and rabbit immunoglobulin (Jackson Immunoresearch, Cat#: 011-000-003 and 006-000-002) in FACS buffer (PBS containing 1% FBS and 0.01% sodium azide) or Fc-receptor block (Innovex, Cat#: 50-486-808). Cells were labeled with an antibody panel targeting CD1b expression (Table S2). Viability of cells was assessed via Zombie Yellow amine-reactive permeability dye (BioLegend, Cat# 423103). Data were acquired using a spectral unmixing Cytek Aurora flow cytometer (Cytek, Fremont, CA) and analyzed via Flowjo software version 10.8 (Flowjo, Ashland, OR) using a minimum of 100,000 events (Fig. S3). CD1b expression was inferred by comparing the mean fluorescence intensity (MFI) values for lung and spleen cells from each infected guinea pig to naïve guinea pigs after applying the same gating strategy.

### CD1 cytotoxicity

Building on prior evidence for cytotoxicity with lipid antigens (29, 30), we measured the cytotoxic activity of CD1b-restricted immunity over the course of Mtb infection. CD1b fibroblast transfectants served as target cells and were incubated with non-adherent cells isolated from the lung or spleen of each animal. Detection of cytotoxicity, assayed as membrane permeability, was performed using a published method (31). In brief, 1×10^7^ CD1b1 and CD1b3 transfected cells were collected and incubated with 0.5 μM Cell Trace Violet (Thermofisher, C34557) for 15 minutes in RPMI media. After quenching with 10 ml of RPMI containing 10% FBS for 10 minutes, cells were washed twice and plated in a 96-well plate at 5×10^4^ per well. Free lipid solutions were prepared as for the intradermal skin testing and resuspended in RPMI media using mock-loaded or unloaded controls for determination of lipid-antigen specific cytotoxicity. Cells and lipid were incubated overnight to allow for lipid cellular trafficking and lipid presentation in the context of CD1b. These target cells were then incubated with non-adherent inflammatory cells derived from the lung and spleen from either naïve or Mtb-infected guinea pigs for 4 hours. Incubated cells were then stained for viability using Zombie Yellow dye, diluted at 1:500 in PBS dilution then added at 100μl per sample (BioLegend, Cat# 423103). Data were acquired using a Cytek Aurora spectral unmixing flow cytometer (Cytek, Fremont, CA) and analyzed via Flowjo software version 10.8 (Flowjo, Ashland, OR) using a minimum of 50,000 events with fibroblasts identified based on size and presence of CellTrace Violet signal (Fig. S4). Mock-transfected fibroblasts, containing the empty vector with no CD1b construct, or CD1b1 and CD1b3 transfected fibroblasts without added lipid antigen were used as negative controls. Data were normalized to background using the mean cellular death among CD1b1 and CD1b3 transfectants without addition of effector cells (1:0 target to effector ratio).

### Kinetic expression of specific CD1b orthologs by qRT-PCR relative gene expression

Kinetic gene expression of CD1b was separately measured with ortholog-specific primers (CD1b1-b4) designed using Geneious software (Auckland, NZ) via a 4-sequence alignment and selecting for non-consensus primer sets, which were then synthesized by Integrated DNA Technologies (IDT) (Coralville, IA) (Table S3). Primer set specificity was evaluated via the use of synthetic gene segments from IDT for each CD1b sequence (Fig. S5), demonstrating specificity for each of the CD1b1-4 orthologs without cross-reactive detection (Fig. S6). For RNA isolation, freshly isolated lung tissue was minced in RNALater (Sigma, cat#: R0901), incubated at 4°C for 24 hours, then frozen at -80°C until RNA extraction. Tissue was thawed at room temperature and approximately 50 mg of tissue homogenized in RLT buffer (Qiagen) containing β-mercaptoethanol using Lysing Matrix A tubes (VWR, cat# 75784-610) in a bead beater instrument at full speed for two cycles of 20 seconds (MP Biomedical, cat# C321001). Proteinase K was added to the homogenate, incubated at 55°C for 15 minutes, then RNA was extracted with the RNeasy mini kit (Qiagen, cat# 74106). Nucleic acid was eluted from the column, digested with DNase-I for 30 minutes at 37°C, then purified again using the RNeasy mini kit. Sample concentration was measured using a Nanodrop spectrophotometer and quality analyzed on an Agilent TapeStation. Generation of cDNA was performed using the iScript cDNA synthesis kit (BioRad cat# 1708890), reverse transcribing 1 μg of RNA per reaction. 25 ng of cDNA template was added to an iScript Sybr Green master mix containing 1.5mM MgCl_2_ and 0.1 μM of each forward and reverse primer. Relative gene expression was measured by normalizing log_2_-fold expression to reference genes HPRT and β-actin using the ΔΔCt method, as previously described (32, 33).

### Distribution of tissue CD1 expression by immunohistochemistry

Tissues from necropsy were embedded in OCT (Thermofisher, cat#: 23-730-571) and cryopreserved in a liquid nitrogen bath, stored at -80°C, and sectioned using a cryotome maintained at -16 to -18°C, and a 7 μm section adhered to charged glass slides. After sectioning, tissues were immediately placed in a 100% chilled methanol fixation for 45 minutes to fix the tissue and inactivate the Mtb organism. All DAB chromogenic and fluorescent immunohistochemistry protocols were completed on methanol fixed tissue sections utilizing a Leica Bond RXm automated slide stainer (Tables S4 and S5). Fluorescent multiplex IHC was achieved using Opal fluorochromes 570 and 690 (Akoya Biosciences, cat# 570- FP1488001KT, 690- FP1495001KT, 690- FP1497001KT) to detect CD1-ortholog specific expression and distribution across myeloid/histiocytic, based on morphology, and specific detection of B cell subsets, based on Pax5 transcription factor expression (clone 24, CellMarque, cat# 312M). All tissues were blocked in 2.5% goat serum solution (Vector Labs, cat# S-1012-50) and primary antibodies diluted in 2.5% goat serum. Primary antibodies were detected using a goat anti-mouse HRP secondary (Vector Labs, cat# MP-7452-50) and incubated with DAB substrate for chromogenic IHC or Opal fluorochromes for fluorescent IHC protocols. Quantification of immunohistochemistry was performed using Visiopharm (Hoersholm, Denmark). A nuclei-detect application was developed to discern between DAPI stained nuclei and Pax5 positive nuclei as well as identifying all cells within tissues, then further delineating cells based on CD1b expression.

### Quantification of tissue bacterial burden

Lung bacterial burden was quantified as previously described (34). In brief, tissue homogenate was plated on 7H11 agar quad plates in 5-fold serial dilutions, incubated for 8 weeks and counted.

### Statistical analysis

All analyses, with the exception of the cytotoxicity assay, were performed utilizing Prism version 9.4.1 (GraphPad, San Diego, CA). Data distribution was evaluated via a Shapiro-Wilks test. Based on distribution, a One-Way ANOVA test or Kruskal-Wallis test was performed to determine statistical significance for flow cytometric analysis, gene expression, and histologic quantifications. Statistical significance was set at p≤0.05. For the cytotoxicity assay, analyses were performed using R Statistical Software (v4.2.1; R Core Team 2021). AUC was calculated based on the ratio of effector to target cells and normalized percent killing via the metrumrg R package (v5.57) (35). A 2-factor ANOVA with Sidak’s correction was used to determine the significance between groups or time with the rstatix R package (v0.7.0) (36).

## RESULTS

### Enumeration of tissue bacterial burden

The burden of Mtb was assessed via culture of lung homogenates and the formation of colony forming units (CFU). CFU counts were consistent with validated and published low-dose aerosol exposure in the guinea pig model (Fig. S7)^27^. Mean CFU per gram of tissue were 4.56 ± 0.30, 4.79 ± 0.45, and 4.76 ± 0.13, for days 14, 30 and 60 of infection, respectively.

### Mtb lipids fail to elicit an intradermal response

Testing of intradermal T cell response to purified protein derivative (PPD) is a mainstay of TB diagnosis that relies on antigen specific recall T cell responses. To assess whether Mtb lipid antigens alone could elicit an immunologic recall response during Mtb infection, infected and naïve guinea pigs were injected with glucose monomycolate or mycolic acid lipid in either a PBS or liposomal formulation. Regardless of formulation administered, synthetic lipid antigens delivered at 14, 30 or 60 days of infection failed to elicit a detectable delayed-type hypersensitivity response to pure antigens. In contrast, and as expected, Mtb PPD elicited a dermal response during at day 30 and day 60 of infection, matching the expected time for early and late adaptive phases of T cell response, with no antigen-specific response to PPD identified at day 14 of infection (Fig. S8).

### CD1b expression during Mtb infection on lung and spleen leukocytes

*In vitro*, human antigen-presenting cells can induce CD1b in response to mycobacterial exposure, suggesting a natural response whereby CD1b can locally function near infection (15). To examine the kinetics *in vivo*, we evaluated the influence of Mtb infection on CD1b expression over the course of TB disease in two independent *in vivo* infection studies in spleen and lung of Mtb-infected guinea pigs among viable CD45+ leukocytes. An increased frequency of CD1b+ cells were demonstrated among infected animals compared to time matched naïve controls via flow cytometric analysis using the 1B12 antibody that recognizes multiple CD1 isoforms (Fig. 1A-D). Inflammatory cells derived from lung parenchyma had similar basal surface CD1b expression during the initial innate response at day 14 of infection, with 12 % ± 6.3% CD1b+ among infected and 10% ± 6.0% among naïve controls. The frequency of CD1b-expressing cells in lung of infected animals was greatest at day 30 of infection, where there was a mean 44% ± 11% CD1b+ cells in infected lung, compared to 10 % ± 6.0% CD1b+ cells among naïve, uninfected guinea pigs. CD1b expression among infected guinea pigs at day 30 of infection was significantly greater than naïve time-matched guinea pigs and kinetically compared to infected guinea pigs at either day 14 or day 60 of infection (p≤0.0001). The frequency of CD1b expressing leukocytes decreased with infection chronicity in the late adaptive immune phase at day 60 of infection, 24% ± 6.3% CD1b+. Guinea pigs at day 60 of infection had a significantly greater frequency of CD1b+ leukocytes compared to animals at day 14 of infection (P≤0.05) but was not significantly greater than time matched naïve guinea pigs (Fig. 1B). Splenic CD45+ leukocytes followed similar surface CD1b expression with peak expression at day 30 (37.8% ± 8.7% CD1b+).

**Figure 1:**
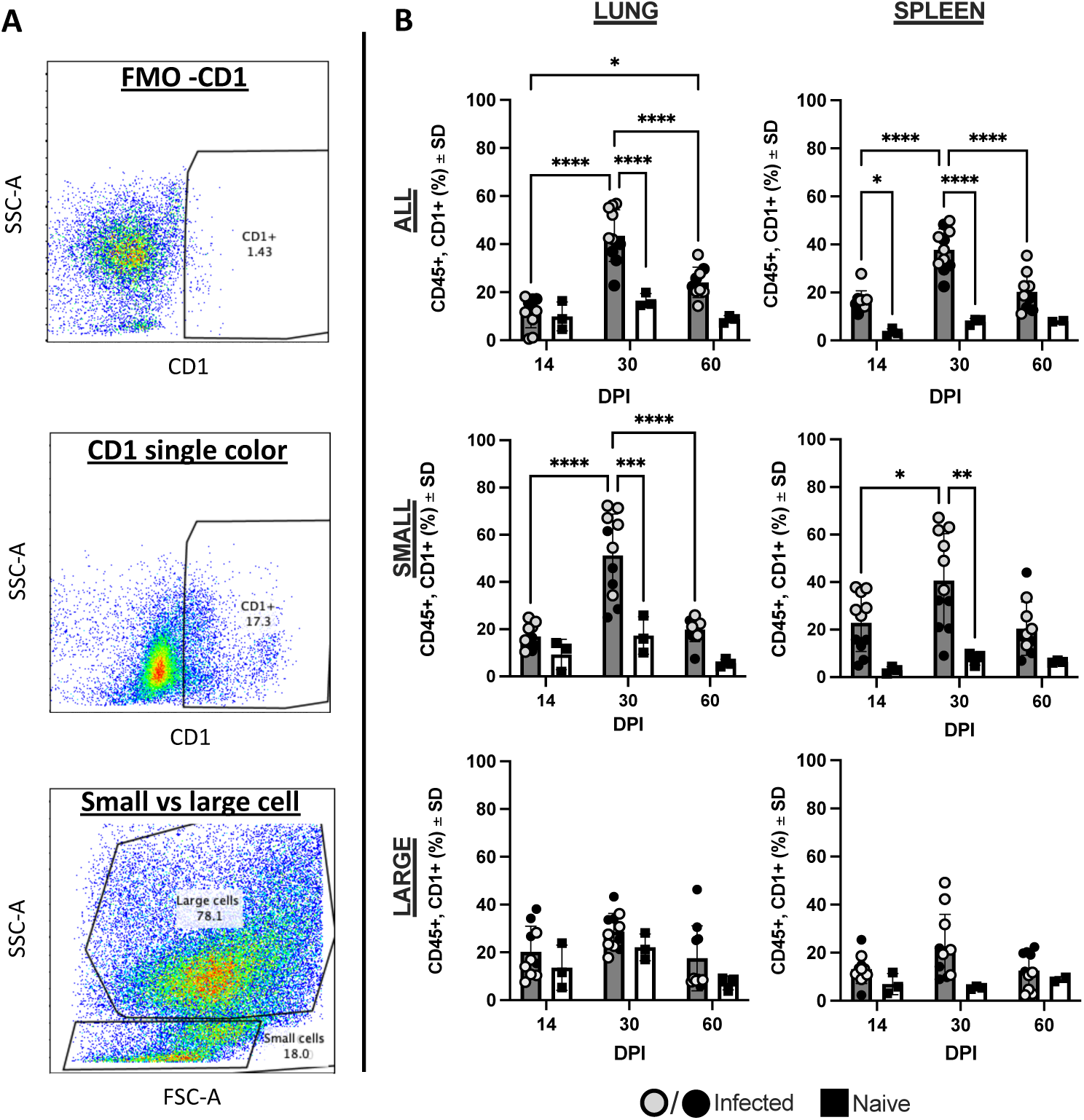
CD1b expression over the course of Mtb infection. Cells were gated for live, CD45+ cells and evaluated for overall CD1 expression using the anti-CD1b monoclonal antibody, 1B12. Open and closed circles represent two separate experimental sets of identical study design. Naïve, time-matched controls were performed solely in one of two experiments. (A) Scatter plot gating of CD1 positive cells, demonstrating fluorescence-minus-one FMO gating control (top), gating of CD1b-expressing cells (middle), and gating strategy for isolating small vs large CD45+ leukocytes (bottom). (B) Proportion of cells expressing CD1b among lung- and spleen-derived cells as all CD45+ cells, delineated small leukocyte populations, and delineated large leukocyte populations. * P≤0.05, **P≤0.01, ***P≤0.001, and **** P≤0.0001; (B) Two-way ANOVA with Tukey post-hoc correction for multiple comparisons.

Owing to the lack of an anti-guinea pig CD3 antibody for flow cytometry, we sought to further discriminate the CD45+ leukocyte populations expressing CD1b over the course of infection based on scatter parameters. Cells were gated by forward and side scatter to differentiate small and large leukocyte populations, previously shown to enrich for lymphocyte and myeloid populations, respectively (27, 28, 37). Large cell leukocytes from both the lung and spleen had low basal surface expression of CD1b with no significant kinetic upregulation over the course of Mtb infection or compared to time-matched naïve control guinea pigs (Fig. 1C). In contrast, lung derived small cell leukocytes had significant upregulation at day 30 of infection (51.2% ± 17.4% CD1b+) compared to naïve guinea pigs (p≤0.001) and exposed guinea pigs at Day 14 (17% ± 5.1% CD1b+), as well as day 60 post-infection (19.9% ± 5.2% CD1b+) (p≤0.0001). Similarly, CD1b expression peaked among splenic small cell leukocytes at day 30 post-infection (40.6% ± 19.7%) (Fig. 1D).

To evaluate the degree of CD1 modulation on individual CD1-positive antigen presenting cells, mean fluorescence intensity (MFI) was measured in Mtb-infected guinea pigs compared to naïve, uninfected animals. In correlation with the cellular breakdown by scatter parameters, MFI was examined in all CD45+ cells, large leukocytes, and small leukocytes (Fig. S9). Increased surface expression of CD1b was observed primarily in lung-derived cells at day 30 of infection compared to naïve guinea pigs, the same endpoint when greater frequency of CD1b+ cells were observed (p ≤0.05). The *in vivo* upregulation was delayed, as no significant differences were observed at 14 DPI. These data suggest that increased expression of CD1b over the course of Mtb infection is the result of both increased cell frequency and upregulation of surface expression on cells capable of expressing CD1b.

### CD1b orthologs have distinct spatial cellular expression profiles

Having identified changes in CD1b protein expression among leukocytes responding to Mtb infection in tissue using a reagent that cannot distinctly discriminate isoforms, we next sought to evaluate the specific CD1b orthologs. The monoclonal antibodies, 1D12 and MsGp9, are specific for CD1b1 and CD1b3, respectively (Fig S1). Lung and spleen tissue from naïve guinea pigs were evaluated for expression of these specific orthologs by immunohistochemistry (Fig 2A-D). Cell- and ortholog-specific expression patterns were evident across lung and spleen tissue. In lung, alveolar and interepithelial cells within the airways were identified with 1D12, which are anatomically and morphologically consistent with alveolar and interepithelial macrophages. In contrast, MsGp9 stained the B cell regions of the splenic white pulp (Fig S10), corroborating previous literature (20). In contrast to the MsGP9 staining, the CD1b1 target of 1D12 within spleen is rare and limited predominantly to cells within the sinuses of the splenic red pulp.

**Figure 2:**
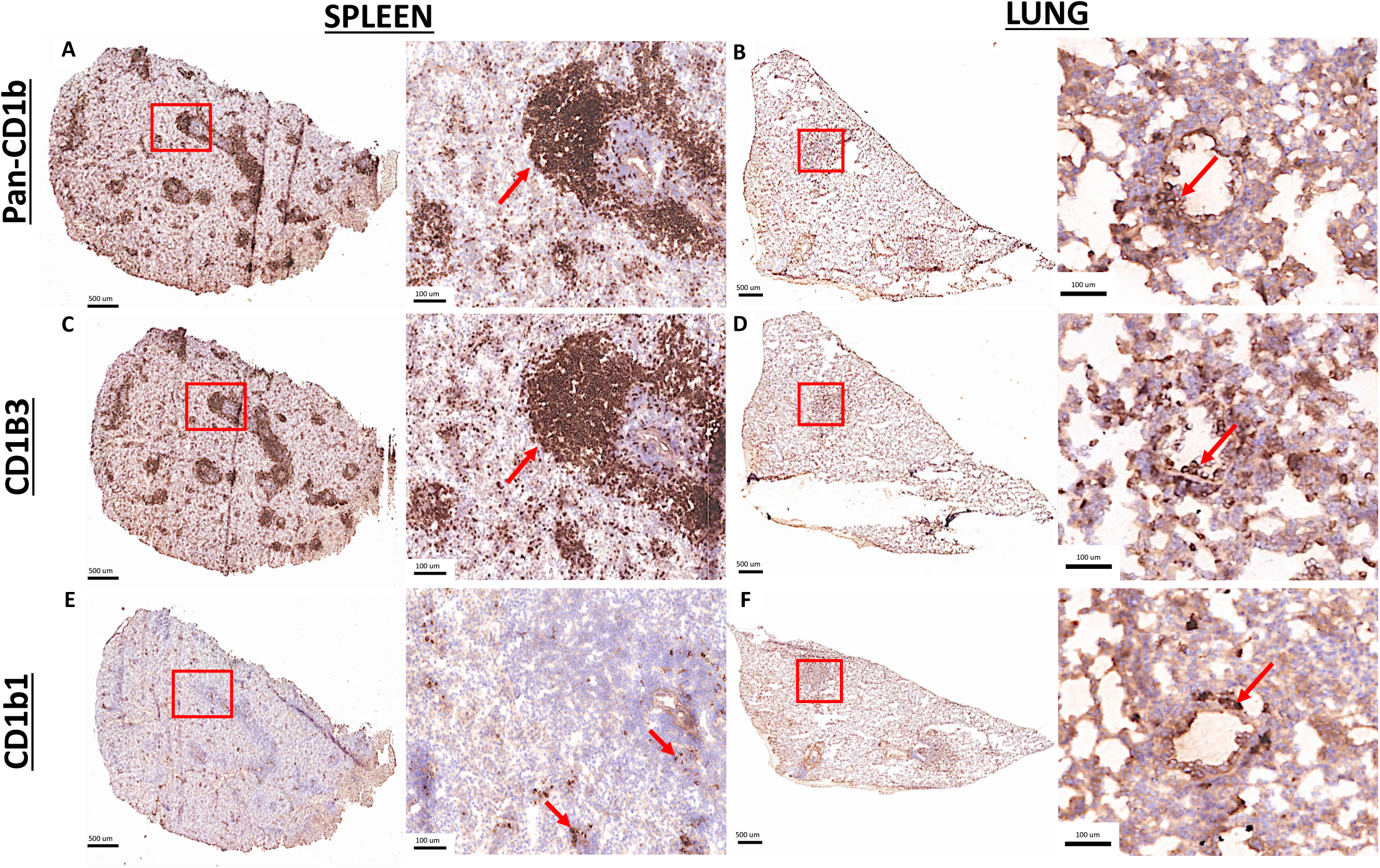
Tissue distribution of specific CD1b orthologs. Fresh-frozen naïve lung and spleen tissue cryosections were fixed in absolute methanol and labeled by immunohistochemistry. (A-B) Distribution of all CD1b orthologs with pan anti-CD1b antibody, 6B5. (C-D) Distribution of the CD1b3 ortholog detected with the anti-CD1b3 monoclonal antibody, MsGP9. (E-F) Distribution of the CD1b1 ortholog detected with anti-CD1b1 monoclonal antibody, 1D12. High magnification images are representative of the areas delineated by red boxes in each figure A-F. Red arrows point to cells immunolabeled by each CD1-targeting monoclonal antibody.

The same chromogenic IHC approach and antibodies were utilized to evaluate CD1b ortholog spatial expression within lung granulomas. Serial sections of lung from an infected guinea pig at the day 30 endpoint were evaluated. IHC performed with the pan-CD1b antibody, 6B5, highlights a population expressing CD1b within the lymphocytic regions at the periphery of primary granulomas. Further delineating ortholog-specific expression, IHC with the CD1b3-specific MsGp9 clone demonstrated strong membranous immunolabeling among morphologically small cells consistent with lymphocytes (Fig. 3A**, B)**. In contrast, CD1b1 assessed by IHC with clone 1D12, showed weak membranous immunolabeling in the wall of the granuloma among the region of cell infiltrate also expressing CD1b3 (Fig 3C). The clear differences across tissue and cellular distribution of CD1b1 and CD1b3 expression suggest that tissue compartments and cellular populations may have specific regulation of CD1b orthologs in naïve tissue and during the course of Mtb infection.

**Figure 3:**
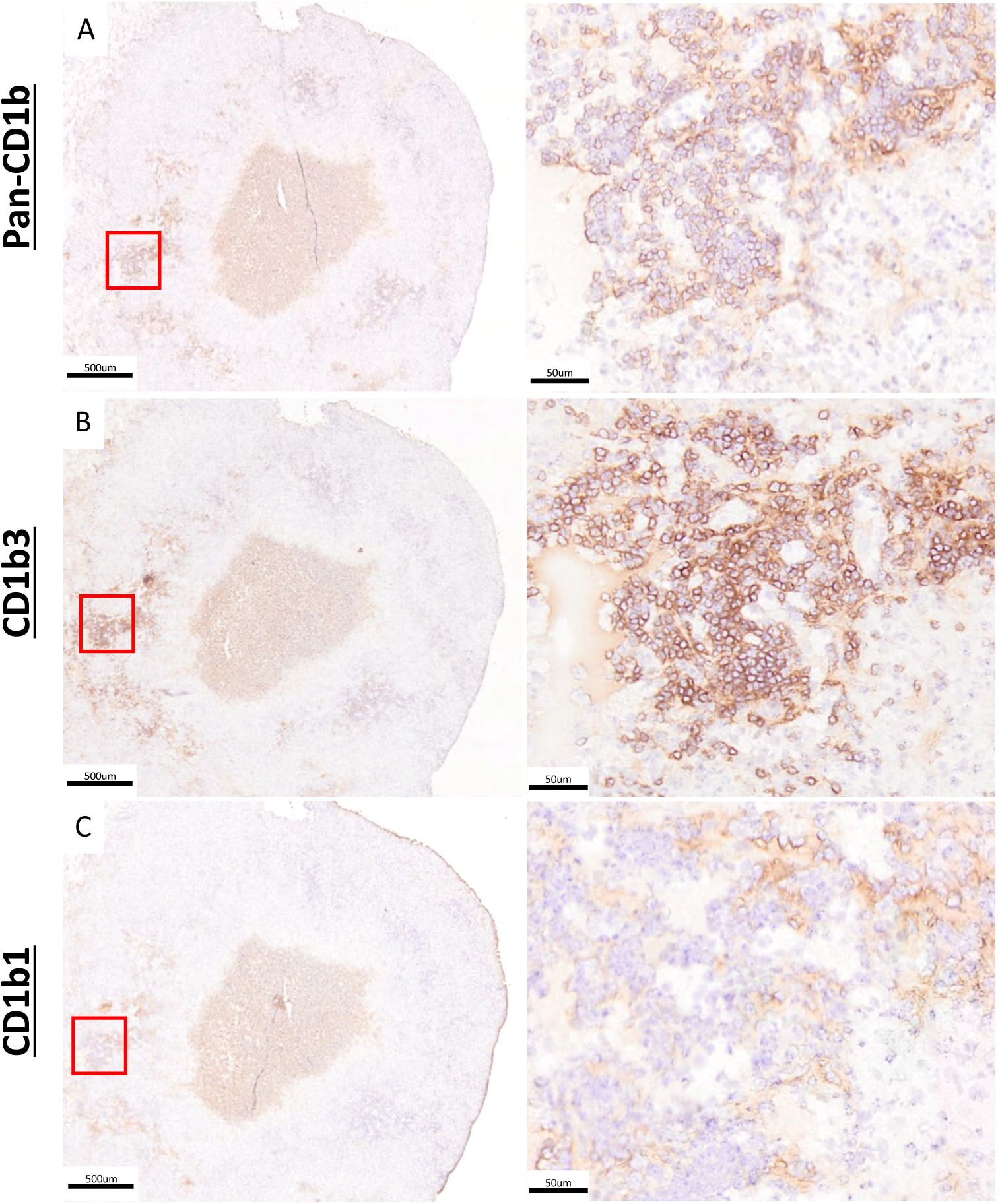
Cellular distribution of CD1b expression in the pulmonary granuloma. Fresh frozen serial sections of a lung granuloma from an Mtb-infected guinea pig at 30-days post-infection were fixed in absolute methanol and labeled by immunohistochemistry using monoclonal antibodies against all CD1b orthologs, or specifically targeting CD1b1 and CD1b3. (A) Distribution of all CD1b orthologs detected with 6B5; (B) Distribution of the CD1b3 ortholog detected with MsGP9. (C) Distribution of the CD1b1 ortholog detected with 1D12. High magnification images are representative of the areas delineated by red boxes in each figure A-C.

Prior studies in humans indicated that CD1b upregulation is mediated mainly through transcriptional control (15). As a second approach to dissect how specific CD1b orthologs are regulated during the course of Mtb infection, we designed ortholog-specific primers targeting CD1b1-b4 and evaluated kinetic changes in relative gene expression in total RNA isolated from infected and naïve lung. Relative gene expression of CD1b orthologs (CD1b1-b4) performed on total lung cDNA demonstrated altered expression of CD1b in infected guinea pigs compared to total lung cDNA from naïve guinea pigs (Fig. 4). Collectively, all CD1b orthologs increased expression in the course of Mtb infection, with peak expression for CD1b1-3 occurring at day 30 of infection, consistent with results achieved by flow cytometry and successfully capturing the kinetic expression over the course of innate and adaptive immunity. CD1b2 showed the greatest increase in relative expression during infection, beginning at day 14 and reaching a peak at day 30 with a 36-fold mean increase (147.3 ± 86.1; p≤0.01) compared to naïve controls. In contrast, CD1b4 peaked early at day 14 of infection with an 18-fold mean increase (73.9 ± 55.84; p≤0.01) then progressively waned over days 30 and 60 of infection. Thus, the timing of CD1b transcriptional induction and waning is similar across CD1b orthologs, although antibody-based detection shows expression of different cell types.

**Figure 4:**
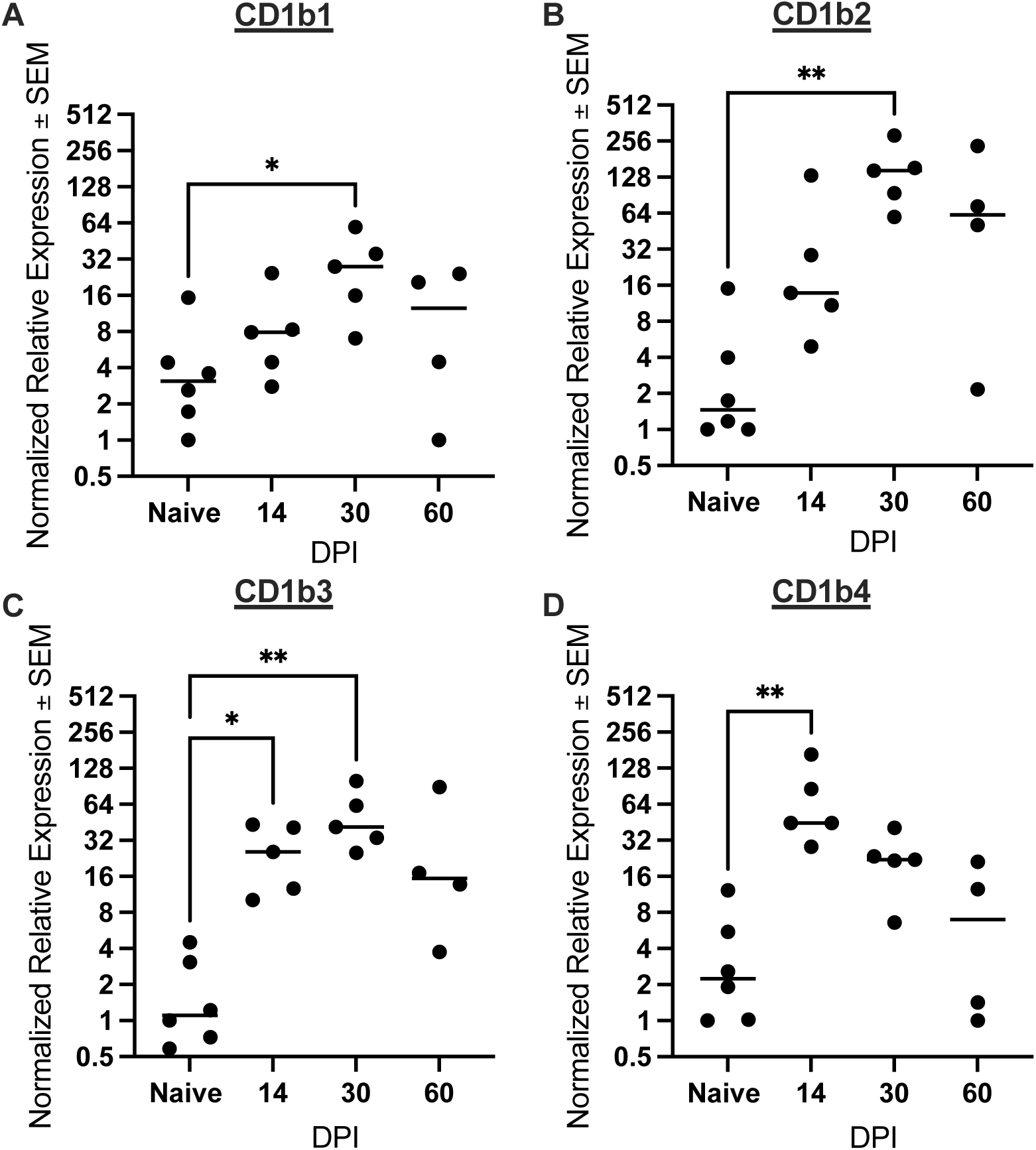
Kinetic gene expression of each CD1b ortholog over the course of Mtb infection. Relative gene expression of each ortholog was measured by quantitative RT-PCR on total RNA isolated from lung samples of Mtb-infected and naïve uninfected guinea pigs. Expression was normalized to the naïve control animal having the lowest expression of the CD1b1 ortholog, designated at a relative expression of 1. *P≤0.05; **P≤0.01; One-way ANOVA multiple comparison with a Dunnet correction.

### CD1b expression patterns on lymphocytes in granulomas

In humans, all four CD1 proteins are expressed on thymocytes, CD1c and CD1d are on B lymphocytes, and CD1b expression is usually restricted to myeloid cells (15). Our new data for CD1b on guinea pigs demonstrated two tissue patterns with apparent expression of CD1b on large and small leukocytes, which are likely to be myeloid and lymphoid cells respectively, would be divergent from the human system. Specifically, increased frequency of CD1b-expression on small CD45+ cells via flow cytometric analysis, and chromogenic IHC highlighting small cells morphologically consistent with lymphocyte expression patterns not seen in humans, so we further pursued CD1b expression via multiplex IHC to identify the cell phenotypes and locations of CD1b-expressing leukocytes within infected lung parenchyma. Multiplex immunohistochemistry showed the majority of CD1b expression is centered on Pax5+ cells located along the periphery of Mtb granulomas, which is phenotypically consistent with B cells (Fig. 5A-D). Correlating with flow cytometric analyses, CD1b was kinetically regulated during the course of Mtb infections, with the highest number CD1b expressing cells appreciated at 30 days post-infection, with a mean 21,369 ± 9,829 CD1+ cells per animal among lung granulomas (Fig. 5E-F).

**Figure 5:**
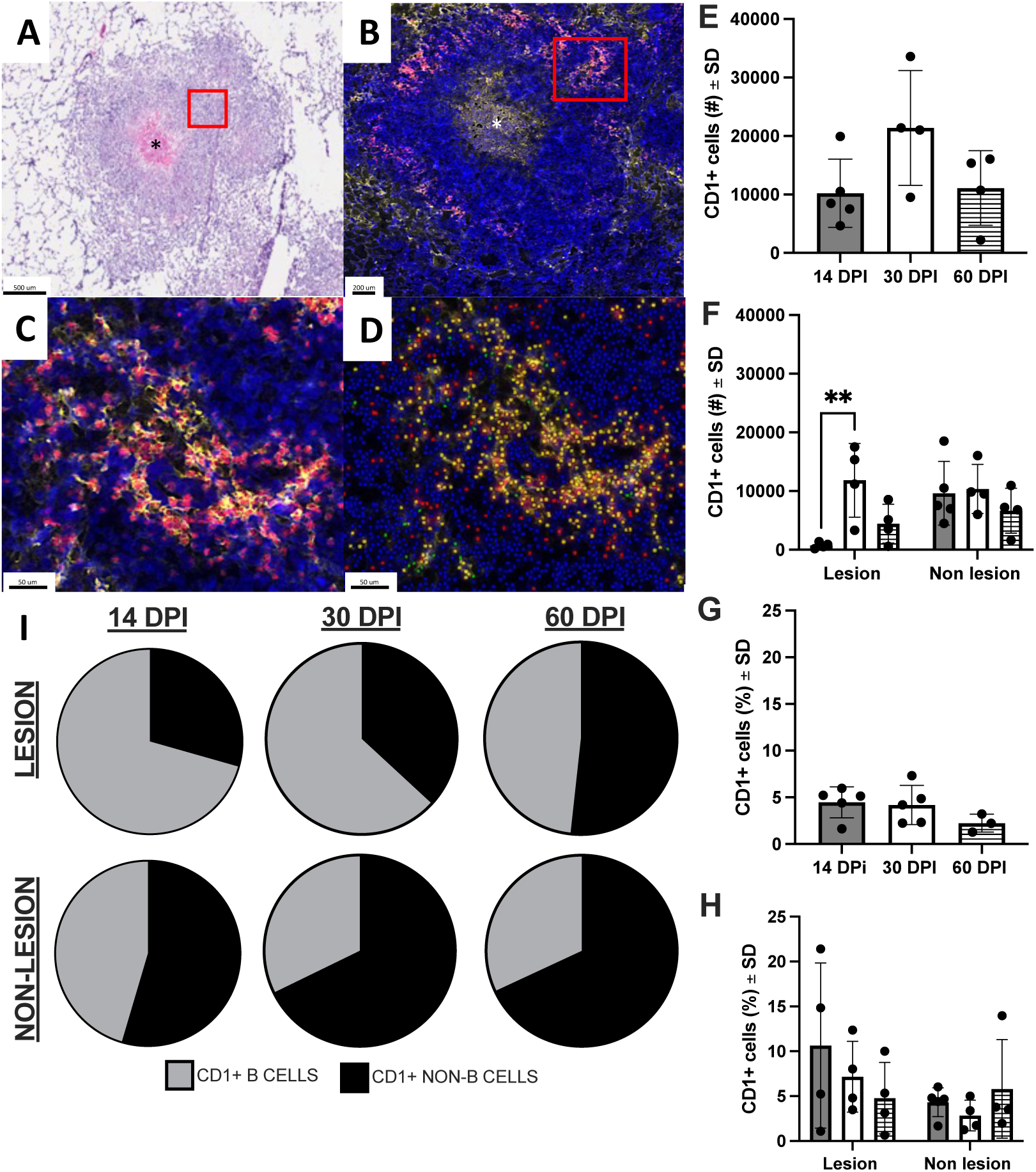
Location and cellular distribution of CD1b expression over the course of Mtb infection. Fresh frozen sections of lung and spleen were fixed in absolute methanol and labeled with antibodies targeting all CD1b orthologs (6B5, Opal 520) or the B cell transcription factor, Pax5 (clone 24, Opal 690), using Opal fluorochrome immunohistochemistry and nuclear counterstain, DAPI. (A-D) Representative sections of a lung granuloma demonstrating: H&E-stained image of the granuloma labeled by IHC (A); Multiplexed IHC detecting CD1b (yellow) and Pax5 (red) demonstrating the distribution of expression across the granuloma (B), with the region of central necrosis marked an asterisk; high magnification view of CD1+, Pax5+ region designated by the red boxes in panels A and B (C); and the same high magnification field of view overlayed with Visiopharm analysis annotation of CD1+ and Pax5+ B cells (yellow), CD1- and Pax5+ B cells (red), or CD1+ and Pax5-representing CD1-expressing cells other than B cells (green) (D). (E-F) Quantitative output of CD1+ cell numbers achieved with Visiopharm image analysis across the entire lung section (E) or separated into lesional and non-lesional regions of each lung section at 14 (grey bar), 30 (white bar), and 60 (hashed bar) days post infection (DPI) (F). (G-H) Proportion of CD1+ cells among all detected cells in whole lung (G) or separated into lesional and non-lesional regions of each lung section at 14 (grey bar), 30 (white bar), and 60 (hashed bar) days post infection (H). (I) Visual designation of CD1+ cell phenotypes as pie diagrams across lesional and non-lesional lung tissue. **P<0.01; one-way ANOVA with a Tukey correction.

Whereas prior studies of mycobacterial modulation of human CD1b expression were mainly conducted on pure cells exposed to mycobacteria or mycobacterial products *in vitro*, these *in vivo* studies provided a window into evolving *in vivo* responses. The concordance of transcripts and protein kinetics was consistent with transcriptional regulation of CD1b proteins. Interestingly, the frequency of CD1+ cells in TB granuloma lesions reflects a large proportion of the initial cells recruited in the early response to Mtb infection, as evidenced by the increased proportion of CD1+ cells among all granuloma-associated infiltrate at day 14 of infection (Fig. 5G-H). As such these data fulfill predictions that mycobacteria induce CD1b needed for presentation of mycobacterial lipids (38), and are consistent with a natural role of CD1b in host response. Because these cells are largely Pax5+ (Fig. 5I), this represents a population of early responding B cells that may represent a gateway to subsequent lipid-restricted immunity. Collectively, evaluation of CD1b expression by IHC methods demonstrates that Pax5+ populations dominate among CD1b-expressing cells in the granuloma, with a surprising paucity of CD1b expression among leukocytes in the walls of TB granulomas that lack expression of the Pax5 marker.

### Infection elicits CD1b-restricted cytotoxicity

To examine the functional response of CD1b-restricted immunity to Mtb lipids over time during infection, non-adherent effector cells isolated from spleen and lung parenchyma were incubated with CD1b1- and CD1b3-transfected fibroblast cell lines loaded with synthetic lipid antigens, mycolic acid or glucose monomycolate. Lung effector cells (Fig. 6A) failed to demonstrate significant antigen-specific cytotoxicity at day 14 of infection, which corresponds to peak innate immunity, though antigen-loaded samples trended toward increased cytotoxicity compared to unloaded controls. However, at time frames exceeding the typical acquired T cell generation, which is about three weeks, cytotoxic effect with lung-derived effector cells were observed at 30 days post-infection, within all conditions except for CD1b3 transfectant target cells loaded with mycolic acid. Though all conditions demonstrated a non-significant trend to higher cytotoxicity in antigen-loaded CD1b-expressing target cells, a significant increase in cellular cytotoxicity was only appreciated in glucose monomycolate-loaded CD1b3 target cells (P≤0.01) and mycolic acid-loaded CD1b1 target cells (P≤0.05). The greatest cytotoxic effect was appreciated comparing lipid-loaded target cells to unloaded target controls from infected animals among CD1b1 and CD1b3 targets loaded with glucose monomycolate and CD1b1 targets loaded with mycolic acid. Where lipid antigen-specific and CD1-restricted responses were detected, there is a corresponding increase in antigen-dependent cytotoxicity with increasing ratio of effector:target cells. Proportions of cytotoxicity, indicated as percent cell death, are listed in Table S6. Comparing only infected samples kinetically over time, all conditions at day 30 post-infection, except for mycolic acid-loaded CD1b3 target cells, have significantly greater cytotoxic effect in lung than the same condition at days 14 or 60 post-infection (P≤0.05) (Fig. S11A).

**Figure 6:**
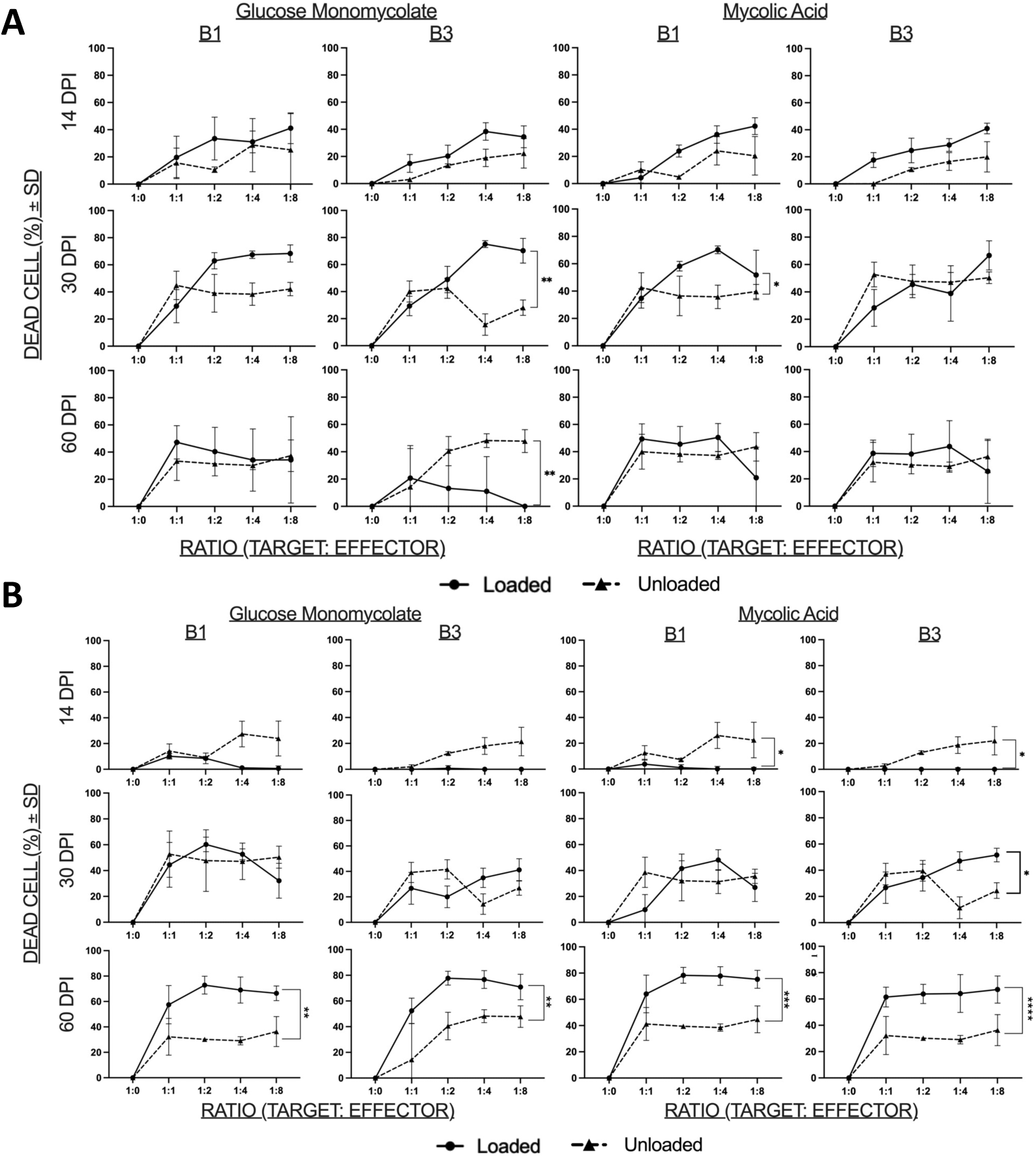
Lipid antigen-specific and CD1b-restricted cytotoxic activity among cells isolated from Mtb-infected lung and spleen. Guinea pig OT1 fibroblasts transfected with CD1b1 or CD1b3 were labeled with CellTrace Violet (CTV) tracking dye then mock-loaded (unloaded) or loaded with synthetic glucose monomycolate or mycolic acid Mtb lipids to be used as target cells in this cytotoxicity assay. Target cell fibroblasts were incubated with non-adherent cells isolated from the lung or spleen of Mtb-infected at indicated ratios for each infection endpoint. Fibroblasts in co-culture were identified by flow cytometric gating on CTV-labeled cells and viability assessed using permeability dye exclusion. A) Fibroblasts incubated with effector cells derived from the lung. B) Fibroblasts incubated with effector cells derived from spleen. All control measures and cytotoxicity comparisons are comprehensively shown in supplemental figures 9 & 10. * P<0.05, **P<0.01, and ***P<0.001; two-way ANOVA with a Sidak’s correction.

Similar to the lung, spleen-derived effector cells (Fig. 6B) lacked significant antigen-dependent cytotoxic function at day 14 for all conditions. At day 30 of infection, the only antigen and CD1b ortholog demonstrating significant cytotoxic activity was seen in mycolic acid loaded CD1b3 target cells compared to unloaded controls (P≤0.05). In contrast to lung, maximal cytotoxic effect with spleen-derived effector cells was observed during chronic disease, at 60 days post-infection, with significant lipid antigen-specific cytotoxicity observed in all conditions, glucose monomycolate-loaded CD1b1 and CD1b3 (P≤0.01), mycolic acid-loaded CD1b1 (P≤0.001) and mycolic acid-loaded CD1b3 (P≤0.0001). Comparing only infected samples kinetically over time, day 30 and 60 endpoints were significantly higher than the day 14 infection endpoint. Additionally, significantly greater cytotoxicity was appreciated in all conditions at 60 days of infection compared to 30 days of infection (Fig. S11B). All comparisons and controls utilized in the lung and spleen cytotoxicity assay, including lipid-loaded and unloaded fibroblasts that do not express CD1, are included in supplemental data (Fig. S12A-B and Table S6).

### CD1b-restricted immunity is not detected among PBMCs

We assessed kinetic CD1b expression and CD1b-restricted cytotoxic capacity among peripheral blood mononuclear cells (PBMCs) to determine how CD1b-restricted immunity is manifested over the course of Mtb infection in peripherally accessible samples. PBMCs, collected via weekly antemortem blood draws and from our specified terminal endpoints, were analyzed for CD1b expression and CD1b-restricted cytotoxicity. Using the 1B12 monoclonal antibody we failed to detect significant upregulation in the PBMC populations by flow cytometry across any of the specified collection points over the course of Mtb infection, or when compared to specified time-matched naïve guinea pigs (Fig. 7A). Due to the requirement for high cell numbers, functional cytotoxicity assays were performed only on PBMCs collected at the specified termination endpoints. Regardless of the infection endpoint, CD1b ortholog-expressing target cell, or Mtb lipid antigen, no cytotoxic effect was demonstrated among PBMCs isolated from infected guinea pigs compared to PBMCs from naïve guinea pigs (Fig. 7B).

**Figure 7:**
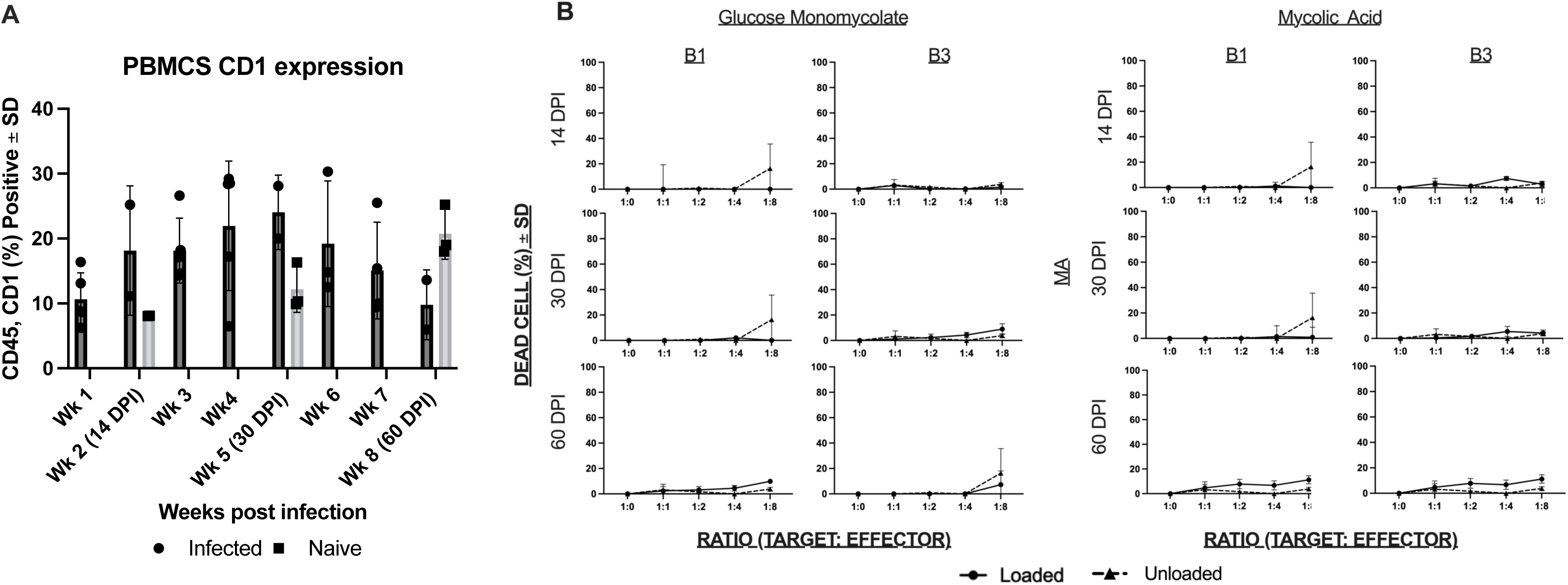
CD1b expression and cytotoxic activity in peripheral blood of naïve and Mtb-infected guinea pigs. (A) Cells were gated for live, CD45+ cells and evaluated for overall CD1b expression using the anti-CD1b monoclonal antibody, 1B12. (B) Guinea pig OT1 fibroblast cells transfected with CD1b1 or CD1b3 were labeled with CellTrace Violet (CTV) tracking dye then mock-loaded (unloaded) or loaded with synthetic glucose monomycolate or mycolic acid. Target cell fibroblasts were incubated with PBMCs derived from Mtb-infected and naïve guinea pig. Fibroblasts in co-culture were identified by flow cytometric gating on CTV-labeled cells and viability assessed using permeability dye exclusion.

## Discussion

There exists clear evidence for clonal (2, 11, 39) and polyclonal responses of human T cells to mycobacterial mycolate antigens (10) across human TB patient populations (26). However, major questions remain regarding the timing and location of CD1-mediated T cell responses during Mtb infection that can be resolved with a tractable *in vivo* model. With regard to timing, CD1-mediated T cell responses might be rapid and innate in nature, like those of NKT cells. However, there is currently limited evidence for pre-expansion of CD1b-reactive T cells, so T cell responses might require T cell expansion, like acquired MHC-restricted responses that develop over three weeks or longer. A third hybrid model is that Mtb rapidly sheds TLR agonists and other signals to induce group 1 CD1 locally at the site of infection, so that CD1 protein expression is linked in time and location to shed of bacterial lipid antigens (38). This latter concept is supported by *in vitro* studies showing upregulation of group 1 CD1 transcripts and proteins within 24 to 72 hours of infection, bacterial exposure or TLR agonist exposure (15, 40) Other studies show loss of CD1 expression on human myeloid cells *in vitro*, which generally occurs at time frames greater than 6 days, when myeloid cell death ensues (41, 42). Currently, specific mechanisms of downregulation are unknown.

Our *in vivo* model begins to dissect the timing and nature of altered CD1b expression during experimental low dose pulmonary infections. We found clear evidence for local upregulation of CD1b proteins and all four guinea pig CD1b orthologs at the transcriptional level. Moreover, we demonstrate local CD1-restricted cytotoxicity targeting two group 1 CD1b proteins, CD1b1 and CD1b3. We further provide *in vivo* evidence for increased transcription occurring prior to generation of lipid antigen specific responses, all of which support the hybrid model and the likely need for weeks of T cell expansion *in vivo* after exposure to the pathogen. These data support and extend prior *in vitro* work in human systems that observed increases in CD1b expression, where the linkage of the timing and extent of change in transcripts and proteins is consistent with transcriptional upregulation. Further, our *in vivo* kinetic data match findings from one human study showing upregulated CD1b protein expression in Mtb-infected human lung with regard to timing, although expression on specific cell types may differ (43).

Considering the location of T cell response *in vivo*, CD1b-restricted cytotoxic activity was demonstrable in infected lung and spleen tissue. Our early efforts to study intradermal responses to lipids may have failed due to the lack of adjuvants or other formulation issues. However, it is notable that *in vivo* guinea pig CD1b system outcomes seen here are different from previously reported MHC-restricted T cells in the sense that we could not detect recall responses in the peripheral blood or skin. Thus, CD1b-restricted responses may be more localized to the site of infection as compared to the systemic nature of MHC-restricted responses. This outcome stands in contrast to a previous study in BCG immunized guinea pigs where recall responses to skin were demonstrated in response to glucose monomycolate in the absence of Mtb infection (44), potentially explained by the purification process of the antigen, intradermal immunization rather than pulmonary infection, or differences between BCG and Mtb organisms.

It is notable that cytotoxicity was identified as a major readout of localized functional activity of CD1b-restricted T cells, a feature that is also reported in human CD1-restricted T cells isolated from peripheral blood (45). We observed cytotoxic activity in an infection dependent and antigen-restricted manner. Synthetic lipid antigens, mycolic acid and glucose monomycolate, both served as effective antigen targets for the induction of cytotoxic activity, which was observed to a greater degree in antigen loaded CD1b1 or CD1b3 transfected fibroblast target cells compared to unloaded controls. Synthetic lipid antigens provide a significant strength as they lack contaminating protein antigens, which may be present in lipid antigens purified directly from the Mtb organism. This approach eliminates the possibility that the antigen-restricted functional response observed in this study is the result of contaminating protein antigens. Altogether, this study provides evidence that CD1-restricted killing of target cells in active TB requires a lipid antigen in association with CD1.

One unexpected difference between the human and guinea pig CD1b systems is that we have identified that group 1 CD1 expression is primarily localized to B cells within granulomatous lesions of the spleen and lung, suggesting this cell population may contribute a previously unrecognized role to the development and maintenance of CD1-restricted immunity. This outcome is different from the human system, where CD1c and CD1d are expressed on B cells, and CD1b is not (46, 47). It was noted that the predominant lesion-associated CD1- expressing cell is the Pax5+ B cell. Although many Pax5+ B cells express CD1b, there is also a substantial population of Pax5+ B cells that are negative for CD1b in granuloma lesions. The difference in function between these cells and their contribution to ongoing granuloma-associated immune responses remains unexplored. We also observed that there is a proportionally greater frequency of CD1+ and Pax5+ B cells in the early immune response at day 14 of infection. This early manifestation of CD1 expression might suggest an efficient and early presentation of lipid antigens and an unexpected role for B cells in the early response to Mtb infection. Of note, there is a paucity of CD1b expression among non-B cell phenotypes within the granuloma, suggesting that granuloma-associated macrophages do not express CD1b orthologs. The cause and consequence of this apparently absent expression among lesion-associated myeloid cells remains unknown. The antibodies used in this study are specific for CD1b, thus, we cannot rule out a different cellular distribution expression profile of CD1c orthologs.

We also observed that CD1b1 and CD1b3 have remarkably different spatial cellular expression patterns in naïve tissues of interest. The presence of CD1b1 on alveolar macrophages in lung, as detected by IHC among cells localized within alveolar spaces, and macrophage-like cells in the splenic red pulp would suggest that CD1b1 is a myeloid-cell associated CD1b ortholog. Additional surface marker reagents are needed to better assign myeloid cell-specific expression. In contrast, CD1b3 was highly expressed within the B cell zone of the splenic white pulp, which can be discretely identified by expression of Pax5 as a highly specific marker of B cell populations. This finding indicates that CD1b3 is a B cell-associated molecule expressed in both naïve and infected conditions. Collectively, these data suggest that CD1b orthologs may serve multiple roles in the contribution to immunity against *M. tuberculosis* infection.

Having now established the presence of CD1-restricted immunity at the primary sites of Mtb infection achieved by low-dose aerosol exposure in the guinea pig model, it remains critical to better understand the contributions of this branch of immunity to TB disease outcome. It is still unknown whether CD1-restricted responses provide a benefit or detriment to the host, and how these antigen-presentation molecules contribute among the mix of a significantly larger repertoire of protein-based antigens in the context of classical MHC class I or class II antigen presentation.

## Funding

NIH 3R01AI049313 to DBM

NIH U19AI111224 to DBM, RJB, and BKP

NIH K01OD016997 to BKP

NIH NIAID 5T32OD010437-19 to MCH and BKP

NIH NCATS Colorado CTSA TL1TR002533 to MCH and BKP

NIH 1310OD030263 S10 Instrumentation Award

## Acknowledgments

The authors wish to acknowledge Dr. Steven Porcelli for provision of guinea pig specific reagents, including pan-CD1b and ortholog specific monoclonal antibodies as well as CD1b transfected OT1 fibroblast cell lines. We thank Dr. Adriaan Minnaard for synthetic mycolic acid and glucose monomycolate antigens.

## References

1. Porcelli S, Brenner MB, Greenstein JL, Balk SP, Terhorst C, Bleicher PA. 1989. Recognition of cluster of differentiation 1 antigens by human CD4-CD8-cytolytic T lymphocyte, vol 341, p 447–450.

2. Beckman EM, Porcelli SA, Morita CT, Behar SM, Furlong ST, Brenner MB. 1994. Recognition of a lipid antigen by CD1-restricted αβ+ T cells, vol 372, p 691–694. Nature Publishing Group.

3. Brigl M, Brenner MB. 2004. CD1: antigen presentation and T cell function, vol 22, p 817–890. Annu Rev Immunol.

4. Brossay L, Kronenberg M. 1999. Highly conserved antigen-presenting function of CD1d molecules, vol 50, p 146–151. Springer Verlag.

5. Chancellor A, Gadola SD, Mansour S. 2018. The versatility of the CD1 lipid antigen presentation pathway, vol 154, p 196–203. Immunology.

6. Dougan SK, Kaser A, Blumberg RS. 2007. CD1 expression on antigen-presenting cells.

7. Pereira CS, Macedo MF. 2016. CD1-restricted T cells at the crossroad of innate and adaptive immunity.

8. Brossay L, Chioda M, Burdin N, Koezuka Y, Casorati G, Dellabona P, Kronenberg M. 1998. CD1d-mediated recognition of an alpha-galactosylceramide by natural killer T cells is highly conserved through mammalian evolution, vol 188, p 1521–1528. J Exp Med.

9. Rossjohn J, Pellicci DG, Patel O, Gapin L, Godfrey DI. 2012. Recognition of CD1d-restricted antigens by natural killer T cells, vol 12, p 845-857. Nature Publishing Group.

10. Gilleron M, Stenger S, Mazorra Z, Wittke F, Mariotti S, Bohmer G, Prandi J, Mori L, Puzo G, De Libero G. 2004. Diacylated sulfoglycolipids are novel mycobacterial antigens stimulating CD1-restricted T cells during infection with Mycobacterium tuberculosis. J Exp Med 199:649–59.

11. Layre E, Collmann A, Bastian M, Mariotti S, Czaplicki J, Prandi J, Mori L, Stenger S, De Libero G, Puzo G, Gilleron M. 2009. Mycolic acids constitute a scaffold for mycobacterial lipid antigens stimulating CD1-restricted T cells. Chem Biol 16:82–92.

12. Moody DB, Guy MR, Grant E, Cheng TY, Brenner MB, Besra GS, Porcelli SA. 2000. Cd1b-Mediated T Cell Recognition of a Glycolipid Antigen Generated from Mycobacterial Lipid and Host Carbohydrate during Infection, vol 192, p 965–976. The Rockefeller University Press.

13. Reijneveld JF, Ocampo TA, Shahine A, Gully BS, Vantourout P, Hayday AC, Rossjohn J, Moody DB, van Rhijn I. 2020. Human γδ T cells recognize CD1b by two distinct mechanisms, vol 117, p 22944–22952. National Academy of Sciences.

14. Montamat-Sicotte DJ, Millington KA, Willcox CR, Hingley-Wilson S, Hackforth S, Innes J, Kon OM, Lammas DA, Minnikin DE, Besra GS, Willcox BE, Lalvani A. 2011. A mycolic acid-specific CD1-restricted T cell population contributes to acute and memory immune responses in human tuberculosis infection, vol 121, p 2493–2503. J Clin Invest.

15. Roura-Mir C, Wang L, Cheng T-Y, Matsunaga I, Dascher CC, Peng SL, Fenton MJ, Kirschning C, Moody DB. 2005. Mycobacterium tuberculosis Regulates CD1 Antigen Presentation Pathways through TLR-2, vol 175, p 1758–1766. American Association of Immunologists.

16. Ulrichs T, Moody DB, Grant E, Kaufmann SHE, Porcelli SA. 2003. T-cell responses to CD1-presented lipid antigens in humans with Mycobacterium tuberculosis infection doi:10.1128/IAI.71.6.3076-3087.2003.

17. Van Rhijn I, Moody DB. 2015. CD1 and mycobacterial lipids activate human T cells doi:10.1111/imr.12253.

18. James CA, Yu KKQ, Gilleron M, Prandi J, Yedulla VR, Moleda ZZ, Diamanti E, Khan M, Aggarwal VK, Reijneveld JF, Reinink P, Lenz S, Emerson RO, Scriba TJ, Souter MNT, Godfrey DI, Pellicci DG, Moody DB, Minnaard AJ, Seshadri C, Van Rhijn I. 2018. CD1b Tetramers Identify T Cells that Recognize Natural and Synthetic Diacylated Sulfoglycolipids from Mycobacterium tuberculosis doi:10.1016/j.chembiol.2018.01.006.

19. Ly D, Kasmar AG, Cheng TY, de Jong A, Huang S, Roy S, Bhatt A, van Summeren RP, Altman JD, Jacobs WR, Adams EJ, Minnaard AJ, Porcelli SA, Moody DB. 2013. CD1c tetramers detect ex vivo T cell responses to processed phosphomycoketide antigens, vol 210, p 729–741. The Rockefeller University Press.

20. Hiromatsu K, Dascher CC, Sugita M, Gingrich-Baker C, Behar SM, LeClair KP, Brenner MB, Porcelli SA. 2002. Characterization of guinea-pig group 1 CD1 proteins doi:10.1046/j.1365-2567.2002.01422.x.

21. Moody DB, Suliman S. 2017. CD1: From Molecules to Diseases.

22. Dascher CC, Hiromatsu K, Xiong X, Morehouse C, Watts G, Liu G, McMurray DN, LeClair KP, Porcelli SA, Brenner MB. 2003. Immunization with a mycobacterial lipid vaccine improves pulmonary pathology in the guinea pig model of tuberculosis.

23. Hiromatsu K, Dascher CC, LeClair KP, Sugita M, Furlong ST, Brenner MB, Porcelli SA. 2002. Induction of CD1-Restricted Immune Responses in Guinea Pigs by Immunization with Mycobacterial Lipid Antigens doi:10.4049/jimmunol.169.1.330.

24. Tahiri N, Fodran P, Jayaraman D, Buter J, Witte MD, Ocampo TA, Moody DB, Van Rhijn I, Minnaard AJ. 2020. Total Synthesis of a Mycolic Acid from Mycobacterium tuberculosis, vol 59, p 7555–7560. Angew Chem Int Ed Engl.

25. Kasmar AG, van Rhijn I, Cheng TY, Turner M, Seshadri C, Schiefner A, Kalathur RC, Annand JW, de Jong A, Shires J, Leon L, Brenner M, Wilson IA, Altman JD, Moody DB. 2011. CD1b tetramers bind αβ T cell receptors to identify a mycobacterial glycolipidreactive T cell repertoire in humans doi:10.1084/jem.20110665.

26. Lopez K, Iwany SK, Suliman S, Reijneveld JF, Ocampo TA, Jimenez J, Calderon R, Lecca L, Murray MB, Moody DB, Van Rhijn I. 2020. CD1b Tetramers Broadly Detect T Cells That Correlate With Mycobacterial Exposure but Not Tuberculosis Disease State doi:10.3389/fimmu.2020.00199.

27. Frenkel JDH, Ackart DF, Todd AK, DiLisio JE, Hoffman S, Tanner S, Kiran D, Murray M, Chicco A, Obregón-Henao A, Podell BK, Basaraba RJ. 2020. Metformin enhances protection in guinea pigs chronically infected with Mycobacterium tuberculosis, vol 10, p 1–11. Nature Publishing Group.

28. Podell BK, Aibana O, Huang CC, DiLisio JE, Harris MC, Ackart DF, Armann K, Grover A, Severe P, Juste MAJ, Dupnik K, Basaraba RJ, Murray MB. 2022. The Impact of Vitamin A Deficiency on Tuberculosis Progression, vol 75, p 2178–2185. Clin Infect Dis.

29. Campos-Martín Y, Gómez del Moral M, Gozalbo-López B, Suela J, Martínez-Naves E. 2004. Expression of human CD1d molecules protects target cells from NK cell-mediated cytolysis, vol 172, p 7297–7305. J Immunol.

30. Lee DJ, Abeyratne A, Carson DA, Corr M. 1998. Induction of an Antigen-specific, CD1-restricted Cytotoxic T Lymphocyte Response In vivo, vol 187, p 433–438. The Rockefeller University Press.

31. Wu X, Zhang Y, Li Y, Schmidt-Wolf IGH. 2021. Improvements in Flow Cytometry-Based Cytotoxicity Assay, vol 99, p 680–688. John Wiley & Sons, Ltd.

32. Ly LH, Russell MI, McMurray DN. 2008. Cytokine profiles in primary and secondary pulmonary granulomas of Guinea pigs with tuberculosis. Am J Respir Cell Mol Biol 38:455–62.

33. Podell BK, Ackart DF, Obregon-Henao A, Eck SP, Henao-Tamayo M, Richardson M, Orme IM, Ordway DJ, Basaraba RJ. 2014. Increased severity of tuberculosis in Guinea pigs with type 2 diabetes: a model of diabetes-tuberculosis comorbidity. Am J Pathol 184:1104–1118.

34. Podell BK, Ackart DF, Kirk NM, Eck SP, Bell C, Basaraba RJ. 2012. Non-Diabetic Hyperglycemia Exacerbates Disease Severity in Mycobacterium tuberculosis Infected Guinea Pigs, vol 7.

35. Bergsma TT, Knebel W, Fisher J, Gillespie WR, Riggs MM, Gibiansky L, Gastonguay MR. 2013. Facilitating pharmacometric workflow with the metrumrg package for R, vol 109, p 77–85. Elsevier.

36. Kassambara A. 2021. rstatix:Pipe-Friendly Framework for Basic Statistical Tests. R package version 0.7.0.

37. Orme IM, Ordway DJ. 2016. Mouse and Guinea Pig Models of Tuberculosis, vol 4. American Society for Microbiology.

38. Moody DB. 2006. TLR gateways to CD1 function. Nat Immunol 7:811–7.

39. Moody DB, Young DC, Cheng TY, Rosat JP, Roura-Mir C, O’Connor PB, Zajonc DM, Walz A, Miller MJ, Levery SB, Wilson IA, Costello CE, Brenner MB. 2004. T cell activation by lipopeptide antigens. Science 303:527–31.

40. Yakimchuk K, Roura-Mir C, Magalhaes KG, de Jong A, Kasmar AG, Granter SR, Budd R, Steere A, Pena-Cruz V, Kirschning C, Cheng TY, Moody DB. 2011. Borrelia burgdorferi infection regulates CD1 expression in human cells and tissues via IL1-beta. Eur J Immunol 41:694–705.

41. Giuliani A, Prete SP, Graziani G, Aquino A, Balduzzi A, Sugita M, Brenner MB, Iona E, Fattorini L, Orefici G, Porcelli SA, Bonmassar E. 2001. Influence of Mycobacterium bovis bacillus Calmette Guérin on in vitro induction of CD1 molecules in human adherent mononuclear cells, vol 69, p 7461–7470. American Society for Microbiology.

42. Stenger S, Niazi KR, Modlin RL. 1998. Down-regulation of CD1 on antigen-presenting cells by infection with Mycobacterium tuberculosis., vol 161.

43. Chancellor A, Tocheva AS, Cave-Ayland C, Tezera L, White A, Al Dulayymi JR, Bridgeman JS, Tews I, Wilson S, Lissin NM, Tebruegge M, Marshall B, Sharpe S, Elliott T, Skylaris CK, Essex JW, Baird MS, Gadola S, Elkington P, Mansour S. 2017. CD1b-restricted GEM T cell responses are modulated by Mycobacterium tuberculosis mycolic acid meromycolate chains. Proc Natl Acad Sci U S A 114:E10956–E10964.

44. Komori T, Nakamura T, Matsunaga I, Morita D, Hattori Y, Kuwata H, Fujiwara N, Hiromatsu K, Harashima H, Sugita M. 2011. A microbial glycolipid functions as a new class of target antigen for delayed-type hypersensitivity. J Biol Chem 286:16800–6.

45. Porcelli S, Morita CT, Brenner MB. 1992. CD1b restricts the response of human CD4-8-T lymphocytes to a microbial antigen. Nature 360:593–7.

46. Allan LL, Stax AM, Zheng DJ, Chung BK, Kozak FK, Tan R, van den Elzen P. 2011. CD1d and CD1c expression in human B cells is regulated by activation and retinoic acid receptor signaling. J Immunol 186:5261–72.

47. Delia D, Cattoretti G, Polli N, Fontanella E, Aiello A, Giardini R, Rilke F, Della Porta G. 1988. CD1c but neither CD1a nor CD1b molecules are expressed on normal, activated, and malignant human B cells: identification of a new B-cell subset. Blood 72:241–7.

